# Pioneer and nonpioneer factor cooperation drives lineage specific chromatin opening

**DOI:** 10.1101/472647

**Authors:** Alexandre Mayran, Kevin Sochodolsky, Konstantin Khetchoumian, Juliette Harris, Yves Gauthier, Amandine Bemmo, Aurelio Balsalobre, Jacques Drouin

## Abstract

Pioneer transcription factors are coined as having the unique property of “opening closed chromatin sites” for implementation of cell fates. We previously showed that the pioneer Pax7 specifies melanotrope cells through deployment of an enhancer repertoire: this allows binding of Tpit, a nonpioneer factor that determines the related lineages of melanotropes and corticotropes. Here, we investigated the relation between these two factors in the pioneer mechanism. Cell-specific gene expression and chromatin landscapes were defined by scRNAseq and chromatin accessibility profiling. We found that in vivo deployment of the melanotrope enhancer repertoire and chromatin opening requires both *Pax7* and *Tpit*. In cells, binding of heterochromatin targets by Pax7 is independent of Tpit but Pax7-dependent chromatin opening requires Tpit. The present work shows that pioneer core properties are limited to the ability to recognize heterochromatin targets and facilitate nonpioneer binding. Chromatin opening per se may be provided through cooperation with nonpioneer factors.

---

In development, cell differentiation leads to establishment of cell identities through the sequential action of transcription factors (TFs) acting as specification or determination factors. These processes involve master TFs that primarily act on the epigenome to open new regulatory chromatin landscapes. This is achieved by a unique class of TF, pioneer factors. Unlike canonical TFs, pioneers bind “closed” chromatin, trigger chromatin opening and thus allow binding of other TFs.

For example, Foxa1 is essential for liver fate and binds regulatory sequences before gene activation^1,2^. Ebf1 also acts as a pioneer factor during B-cell development and Neurod1 and Ascl1 (Mash1) were suggested to function as pioneer factors during neural development^3,4^. Ectopic expression of pioneer factors is sufficient to drive trans-differentiation: thus, C/EBPα can direct trans-differentiation of pre-B cells into macrophages^5^. The most extreme example is the reprogramming of fibroblasts into induced pluripotent stem cells through action of the pioneer factors Oct4, Klf4 and Sox2^6,7^. Similarly, C/EBPα can direct trans-differentiation of pre-B cells into macrophages^5^.

The ability to open chromatin has been implicitly considered as a core property of pioneer TFs and this would allow nonpioneer binding to newly accessible sites. Whether nonpioneers play a role in pioneer-driven chromatin opening has nor been assessed. Here, we used normal and perturbed pituitary differentiation to investigate the pioneer model and establish the specific and/or overlapping functions of the differentiation regulators Pax7 (pioneer) and Tpit (nonpioneer).

Two pituitary lineages express the hormone precursor pro-opiomelanocortin (POMC), the melanotropes and corticotropes. Both require Tpit for POMC expression and cell fate determination^8,9^. Indeed, Tpit implements a secretory cell transcriptional program by activation of scaling factors for translation and secretory organellogenesis^10^ While sharing the secretory POMC identity, corticotropes and melanotropes differ by their functions as these POMC-expressing lineages control corticosteroidogenesis and pigmentation, respectively^11^. The pioneer Pax7 drives melanotrope specification^12^ through deployment of a melanotrope-specific enhancer repertoire^13^.

Here, we first establish that the two POMC lineages share a transcriptional program that is distinct from other pituitary cells and that in addition, they each have a unique program of gene expression. We then show that these two layers of identity (shared and lineage-specific) are reflected at the level of chromatin accessibility and that the shared POMC chromatin landscape requires Tpit. Further, Tpit is required for the opening of the Pax7-dependent melanotrope chromatin landscape indicating that Pax7 and Tpit act together during melanotrope differentiation. Finally, Pax7 and Tpit have complementary roles as only Pax7 can bind heterochromatin while Tpit binding is needed for chromatin opening. In summary, we propose that the essence of pioneer action is the ability to recognize and bind DNA sites in closed chromatin whereas cooperating nonpioneer TFs, such as Tpit, may drive chromatin opening.

## Results

### Single cell analysis reveals the transcriptional diversity of pituitary lineages

The pituitary is a highly specialized organ where each lineage serves as a hormone producing factory. Each cell type is dedicated to the regulation of a specific endocrine organ and responds to specific signals from the hypothalamus and body. We used single cell RNAseq to decipher the transcriptional complexity of the different pituitary lineages. For each lineage, hormone-coding mRNAs are so abundant that they appear as peaks in cDNA libraries (Supplementary Fig. 1A). Profiling of adult mouse male pituitary cells (Supplementary Fig. 1B) was achieved by plotting single cell data using t-distributed stochastic neighbor embedding (t-SNE), a common method^14^ that uses dimensionality reduction to cluster together cells with similar transcriptional profiles (Fig. 1A). This showed that cells expressing the same pituitary hormone cluster together and that they all express Pitx1, a marker of the oral ectoderm origin of the pituitary^11^ (Supplementary Fig. 1C and 1B). We identified 12 clusters composed of endocrine and non-endocrine cells (Fig. 1C). Cluster 1 corresponds to somatotropes as they express the growth hormone (Gh) gene. Lactotropes that produce Prolactin (Prl) are found in cluster 2. Clusters 4 and 5 correspond to melanotropes and corticotropes, respectively; both express the *POMC* gene, yet only melanotropes express *Pcsk2*. Gonadotropes that express the *Lhβ* gene are in cluster 8. We also detected thyrotropes as *Tshβ*-expressing cells; however they did not appear as a separate cluster (Supplementary Fig. 1C). The pituitary stem cells that express Sox2^15^ are found in cluster 7. Although our study aimed at defining the transcriptome of the different pituitary lineages, we also uncovered several non-endocrine cells within the pituitary tissue that do not express Pitx1 (Fig. 1B). We identified endothelial cells (cluster 9), macrophages (cluster 10), posterior pituicytes (cluster 11) and pericytes (cluster 12). Cluster 3 is fragmented in three different groups of cells that express either GH, Prolactin, POMC or Lhβ. We performed differential expression analysis between each sub-cluster 3 and its matching cell type in order to define these subsets. In all cases, cells of these subsets are specifically depleted of ribosomal proteins and enriched for mitochondrial RNA. This was shown to be an artefact of tissue dissociation^16^ and to represent cells affected by the preparation: we excluded cluster 3 from following analyses.

**Fig. 1.**
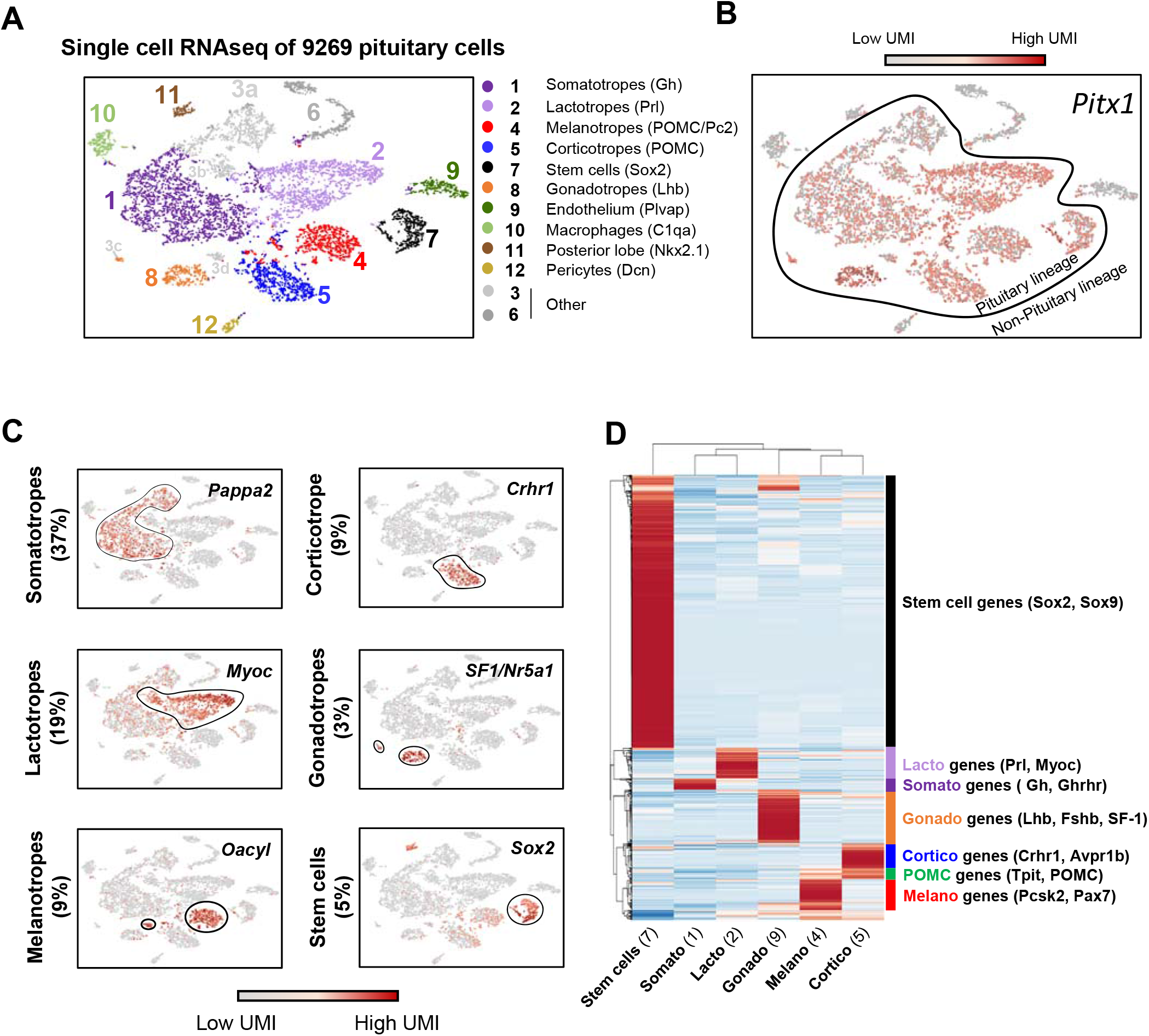
Transcriptional complexity of the pituitary adult gland. A. t-SNE map (t-distributed stochastic neighbor embedding) plot of the 9269 profiled pituitary cells colored by the 12 clusters identified using unsupervised k-means clustering. Cluster identification included expression of the hallmarks gene(s) indicated between parenthesis and the markers shown in Fig. 1C.
B. t-SNE map showing color-coded Pitx1 expression.
C. t-SNE map showing color-coded expression of indicated markers for the major pituitary lineages.
D. Heatmap showing normalized expression of the 1000 most differentially expressed genes (p value<0.05, FC>2 and minimum number of UMI >0.3) in clusters representing the different endocrine and progenitor cells (cluster 1, 2, 4, 5, 7, 8). Rows are centered; unit variance scaling is applied to rows. Both rows and columns are clustered using correlation distance and average linkage.

We then compared the transcriptomes of the different pituitary lineages by performing differential expression analysis between clusters 1, 2, 4, 5, 7 and 8. Genes with two-fold differential expression between each cluster (p value <0.05 and minimum 0.2 UMI in at least one pooled cluster) are shown as a heatmap (Fig. 1D). The Sox2+ stem cell niche is the most transcriptionally divergent from other pituitary lineages based on correlation clustering analyses. The two Pit1-dependent lineages, the lactotropes and somatotropes, are the most different compared to the two POMC lineages and gonadotropes that cluster together. Within the latter group, corticotropes and melanotropes are more transcriptionally correlated together than with gonadotropes. Thus, the two POMC lineages, melanotropes and corticotropes, have both a shared and a specific transcriptional program.

### Lineage-specific chromatin landscapes identify lineage regulators

We next aimed to identify cis-regulatory elements that regulate the transcriptional identity of pituitary lineages. We used ATACseq^17^ to identify the putative regulatory elements accessible in each lineage. We complemented our previously published datasets of purified melanotrope and corticotrope ATACseq with accessibility profiles for gonadotropes and anterior lobe (AL) cells. Gonadotropes were FACS-purified from transgenic pituitaries expressing the LHβ-Cerulean transgene^18^. As control, we isolated the remaining AL cells that are mostly composed of a combination of Pit1-dependent somatotropes and lactotropes. Interestingly, we found that the promoters of hormone genes *POMC, αGSU, Gh* and *Prl* as well as lineage specifiers *Tpit* (*Tbx19*), *Pax7, SF-1* (*Nr5a1*) and *Pit1* show lineage-specific accessibility (Fig. 2A and Supplementary Fig.2A-C). However, the *Pcsk2* promoter is accessible in all pituitary lineages but its numerous distal accessible sites (putative enhancers) are only accessible in melanotropes. Globally, we identified 98926 open chromatin regions across the pituitary lineages (Fig. 2B). Segregation of lineage-specific accessibility yielded 33451 regions opened in all lineages, 14025 regions opened in a combination of three lineages, 20374 in two lineages and finally 31076 opened in only one lineage. Thus, there are regions specifically accessible in melanotropes, corticotropes, gonadotropes or in the AL. In accordance with the close transcriptional correlation between melanotropes and corticotropes, we also found shared regions accessible in both melanotropes and corticotropes (POMC-specific) but closed in gonadotropes and AL. Finally, we identified 13130 pituitary-specific sites that are closed in embryonic stem cells, as well as a set of 20321 regions accessible in both pituitary and ES cells (Supplementary Fig. 2D). These ubiquitous peaks are for the majority (58%) composed of promoter elements (Supplementary Fig. 2E) while regions opened in two or in only one pituitary lineage are mostly distal elements (94% and 96%, respectively). This reinforces the idea that promoter accessibility is established early during differentiation. Further, this suggests that lineage specific opening of promoters tends to be an exception and may be involved in restricting appropriate expression of critical genes such as hormone-coding genes and lineage specifiers.

**Fig. 2.**
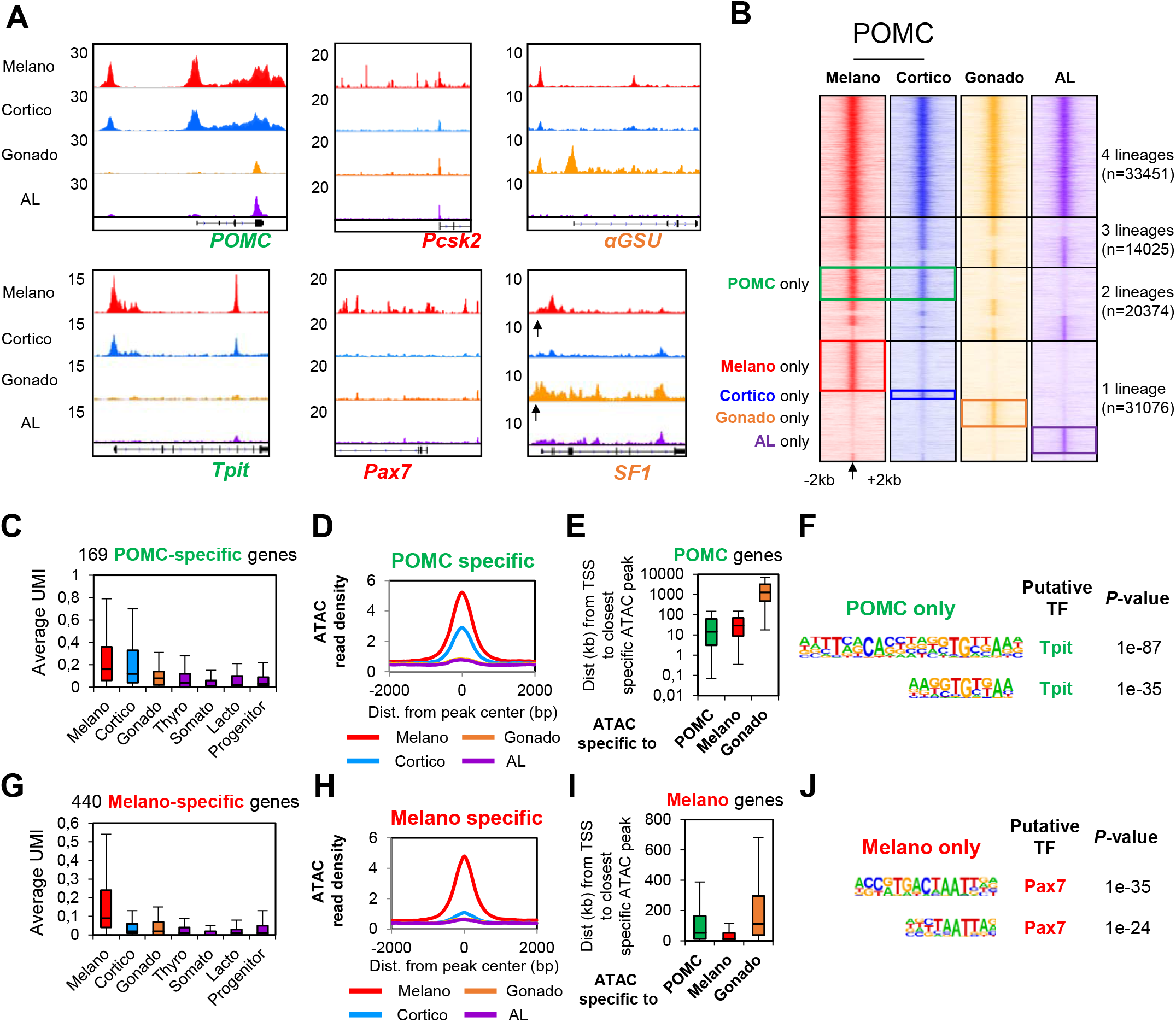
Lineage-specific chromatin access reveals lineage regulators. A. Genome browser view (IGV) of ATACseq profiles at genes marking pituitary lineages: shared POMC markers (green), melanotrope (red) or gonadotrope (orange). The *SF1* promoter is indicated by an arrow.
B. Heatmaps showing ATACseq signals (RPKM) across the different pituitary lineages in a 4kb window around the ATACseq peak center indicated by an arrow. Colored boxes indicate peaks specifically enriched in the indicated lineage.
C. Boxplot showing expression (UMI) of gene markers of the POMC lineages across the different lineages. Center lines show medians; box limits indicate the twenty-fifth and seventy-fifth percentiles; whiskers extend to 1.5 times the interquartile range from the twenty-fifth to seventy-fifth percentiles.
D. Average profiles of ATACseq signals at POMC-specific ATACseq peaks.
E. Box-plot of distances between the TSS of POMC-specific genes and the closest ATACseq peak in POMC, melanotropes and gonadotrope cells. Box plot features as in C.
F. Motif enriched (assessed by HOMER) under POMC-specific ATACseq peaks and not found in other subsets.
G. Box-plot showing expression (UMI) of melanotrope gene markers across the different lineages. Box plot features as in C. H. Average profiles of ATACseq signals at melanotrope specific ATACseq peaks.
H. Box-plot of distances between the TSS of melanotrope-specific genes and the closest ATACseq peak in POMC, melanotrope and gonadotrope cells. Box plot features as in C.
I. Motif enriched (assessed by HOMER) under melanotrope specific ATACseq peaks and not found in other subsets.

Comparison of the spatial relationship between the POMC-specific transcriptional program (Fig. 2C) and lineage-specific accessibility (Fig. 2D) shows that POMC-specific gene promoters tend to be closer to both POMC- and melano-specific open regions (Fig. 2E) and are enriched for the motif of the lineage specifier Tpit (Fig. 2F). Melanotrope genes are also close to their lineage-specific open chromatin regions and their lineage specific chromatin landscape is enriched for the motif of their lineage specifier Pax7 (Fig. 2G-J). Similar relationships were found for the Pit1-dependent lineages, the corticotropes and the gonadotropes (Supplementary Fig. 2F-G).

### Pax7 is required for phenotypical features of melanotropes

The two POMC lineages have shared and specific transcriptional programs and open chromatin landscapes. Their most obvious similarity is expression of the hormone precursor POMC as well as expression of the terminal differentiation factor Tpit. However, POMC expression varies between these two lineages with low (blue) and high expression (red) cells in scRNAseq analyses (Fig. 3A). Indeed, POMC is more highly expressed in Pax7-expressing melanotropes compared to GR (nr3c1)-expressing corticotropes as shown in t-SNE and differential expression Volcano plots (Fig. 3A,B). Analysis of the POMC-EGFP^19^ transgenic pituitaries (Fig. 3C) also showed high and low-expressing cells. Indeed, IL (melanotropes) transgenic pituitary cells have much higher EGFP levels compared to most EGFP-positive corticotrope cells of the AL (Fig. 3D). This indicates that the transgene expressed in both melanotropes and corticotropes is subject to cell-type specific regulation and can be used as a surrogate phenotypical readout. Strikingly in a *Pax7−/−* background, melanotropes express the EGFP transgene at the same level as in AL corticotropes (Fig. 3D, E). Melanotropes are also typically larger and have a more complex organelle content than corticotropes as assessed by forward and side scatter distributions in FACS profiles and this is also lost in Pax7-deficient mice (Supplementary Fig. 3A, B). Thus, all three melanotrope features (high POMC expression, large cell size and granularity) are fully dependent on Pax7 and virtually all melanotropes switch to a corticotrope phenotype in *Pax7−/−* mice (Fig. 3F). However, the proportion of total EGFP-positive cells in AL or IL is not affected by loss of Pax7 (Fig. 3G). Thus, Pax7 implements melanotrope but not the shared POMC cell identity which is under the control of Tpit^9^.

**Fig. 3.**
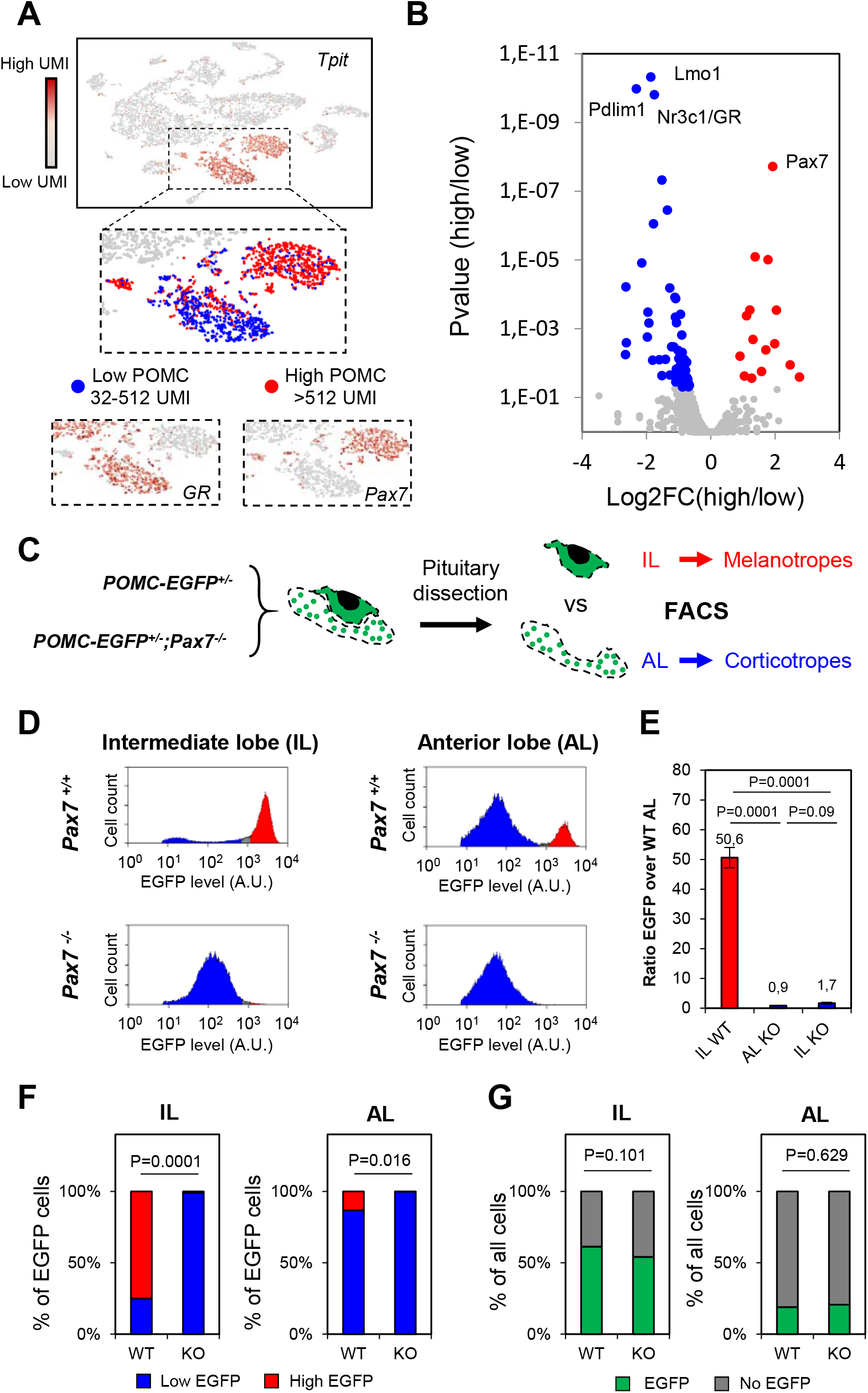
Pax7 implements melanotrope features onto a shared POMC cell identity. A. t-SNE map of adult pituitary cells colored for expression of the POMC lineage regulator Tpit (Tbx19). Enlarged panels showing two clusters of POMC cells with high and low POMC levels, melanotropes expressing Pax7 and corticotropes expressing GR.
B. Volcano plot showing differential transcription factor gene expression (p value vs log2 FC) between the high versus low POMC expressing cells. GR and Pax7 are highlighted.
C. Experimental scheme to assess Pax7 dependence of melanotrope phenotypical features. Transgenic mice expressing POMC-EGFP were crossed into *Pax7−/−* mice and compared to WT. The two pituitary lobes were dissected for each genotype and analyzed by FACS.
D. Representative FACS profiles showing cell populations with different POMC-EGFP transgene levels in intermediate (IL) and anterior lobes (AL) of *WT* (n=5) and *Pax7 KO* (n=3) pituitaries. Blue and red shading labels low and high EGFP levels, respectively.
E. Bar graph showing ratios of EGFP signals in *WT* IL, *Pax7 KO* AL, *Pax7 KO* IL compared to *WT* AL. The analyses included 5 *WT* and 3 *Pax7 KO* replicates. P values were computed using unpaired two-sided t-test.
F. Bar graph showing the proportion of high versus low EGFP expressing cells based on fluorescence signals detected by FACS. P values were computed using unpaired two-sided t-test.
G. Bar graph showing the proportion of EGFP expressing and non-expressing cells based on fluorescence signals detected by FACS. P values were computed using unpaired two-sided t-test.

### Pax7 and Tpit are required for opening cognate lineage-specific enhancer landscape

We next sought to uncouple the specific roles of Pax7 and Tpit for implementation of the shared and melanotrope-specific chromatin landscapes. To do so, we performed ATACseq on IL cells from mice of genetic backgrounds lacking one or both alleles of *Pax7* and/or *Tpit*. We used *Pax7* and *Tpit* double heterozygote littermates as control; these show minimal differences compared to wild-type animals (Supplementary Fig. 4). We found that melanotrope-specific peaks (Fig. 2B) are lost in *Pax7−/−;Tpit+/−* IL whereas they are present in wild-type or double heterozygotes (Fig. 4A) in agreement with previous data^13^ that showed Pax7 requirement for accessibility of melanotrope regulatory modules. Also, we found that Tpit is required for the open status of POMC-specific open chromatin (Fig. 4B). This suggests that Tpit is involved in the pioneering process of the shared POMC lineage enhancers. For example, POMC gene expression critically relies on Tpit and accordingly, both its promoter and enhancer accessibility strongly depend on Tpit but not on Pax7 (Fig. 4C). This is also true for the promoter of POMC-specific gene *Tnxb* and the Tpit-dependent enhancer^10^ of the *Creb3l2* gene (Fig. 4C). Thus, Tpit is required for opening the shared POMC regulatory modules while Pax7 is only required for melanotrope enhancers.

**Fig. 4.**
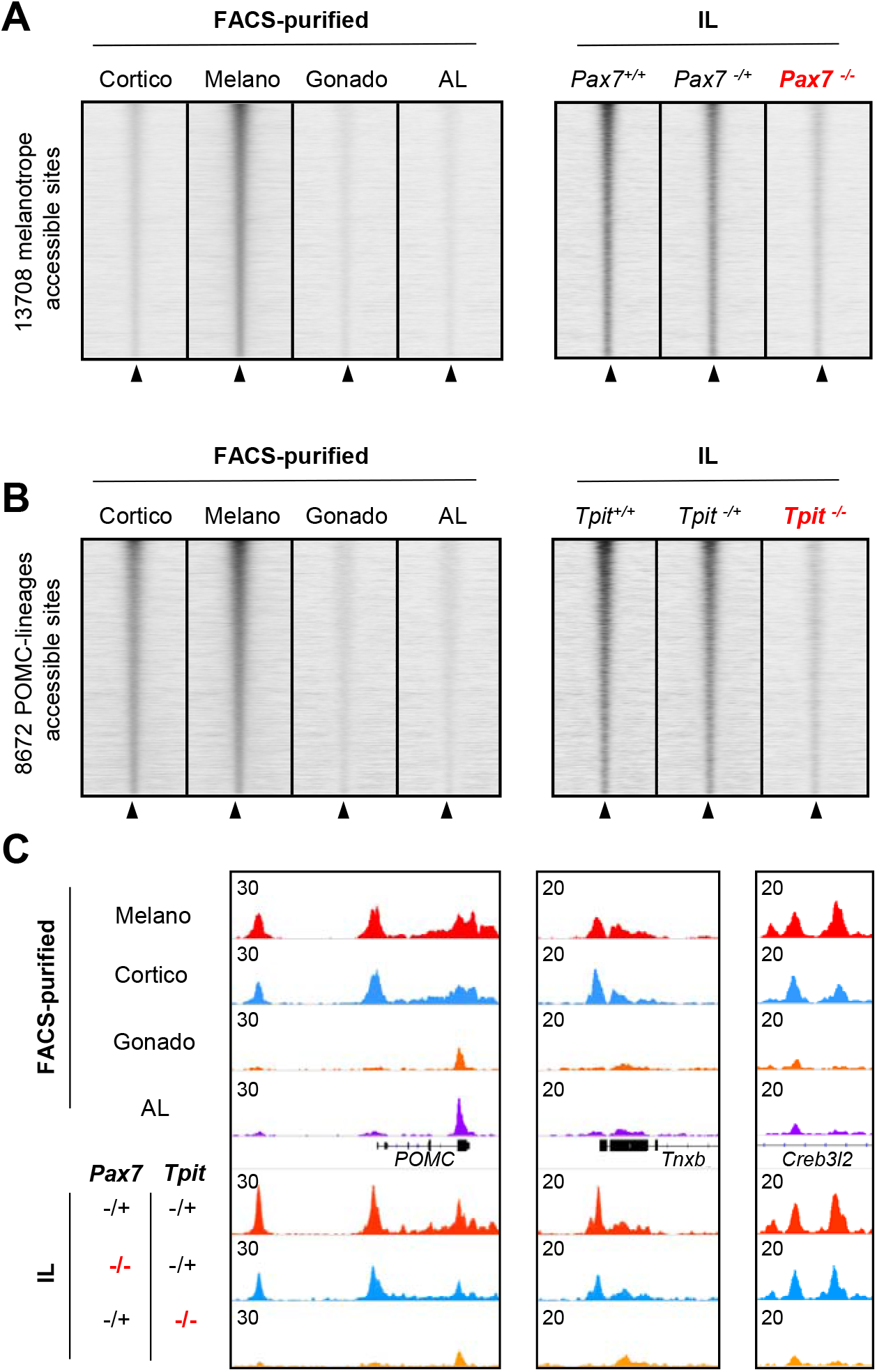
Pax7 and Tpit are required for opening cognate enhancer landscapes. A. Read density heatmaps showing ATACseq signals (RPKM) across the different pituitary lineages in a 4kb window centered at melanotrope specific ATAC peaks (left panel). Right panel shows corresponding ATACseq heatmaps in the ILs of WT, *Pax7^+/−^;Tpit^+/−^* (labeled Pax7^+/−^) and *Pax7^−/−^;Tpit^+/−^* (labeled Pax7^−/−^) mice.
B. Read density heatmaps showing ATACseq signals (RPKM) across the different pituitary lineages in a 4kb window centered at POMC-specific ATAC peaks (left panel). Right panel shows corresponding ATACseq heatmaps in the intermediate lobe of WT, *Pax7^+/−^;Tpit^+/−^* (labeled Pax7^+/−^) and *Pax7^−/−^;Tpit^+/−^* (labeled Pax7^−/−^) mice.
C. Genome browser view (IGV) of ATACseq profiles at POMC-specific ATACseq peaks in the different pituitary lineages and in ILs of *Pax7^+/−^;Tpit+^/−^, Pax7^−/−^;Tpit^+/−^* and *Pax7^+/−^;Tpit^−/−^* mice.

### Tpit is required for Pax7-dependent chromatin opening

In order to define and compare the Pax7 and Tpit-dependent ATACseq landscapes, we performed differential enrichment analysis (p<0.05 and fold changes >2) and found 16024 sites altered in *Tpit−/−* IL (Fig. 5A). Most changes are decreased accessibility (12573 sites) and these show greater quantitative differences compared to sites with increased accessibility (3451 sites). Analysis of these Tpit-dependent sites revealed two subsets, one dependent and one independent of Pax7 (Fig. 5B). In both Tpit-dependent subsets, chromatin accessibility in *Pax7−/−;Tpit−/−* IL is the same as in *Tpit−/−*. In order to ascertain the reliability of these analyses, we validated dependence on Pax7, Tpit or both by qPCR analyses of three independent ATACseq libraries of each genotype (Supplementary Fig. 5). Principal component analysis using all Tpit-regulated sites shows that most of the variance between samples (77%) is explained by component 1 (Fig. 5C). Consistent with the heatmaps of Fig. 5B, wild-type and *Pax7+/−;Tpit+/−* cluster together while *Pax7* knockout are between wild-type and *Tpit* knockout samples. Thus, a subset of Tpit chromatin targets are also dependent on Pax7. We then focused on Pax7-dependent chromatin access to assess whether co-dependency on Tpit and Pax7 is a general feature of Pax7-dependent sites. We found 7057 Pax7-dependent sites (Fig. 5D): most changes are decreased accessibility (6112 sites) and these are greater effects compared to increased sites (945 sites). Strikingly, and unlike Tpit-dependent accessibility, virtually all Pax7-dependent sites are also dependent on Tpit (Fig. 5E). Accordingly, principal component analysis of Pax7-dependent sites showed that Pax7 knockout samples cluster with *Tpit* knockout and Pax7 on component 1 (79% of variance). This indicates that *in vivo*, Tpit is absolutely required for the establishment of the Pax7-dependent chromatin landscape. Thus in both cases, *Pax7* and *Tpit* double knockout samples cluster closely with the *Tpit* single knockout (Fig. 5C, F) showing that loss of both factors does not have a greater impact on accessibility than the loss of Tpit alone.

**Fig. 5.**
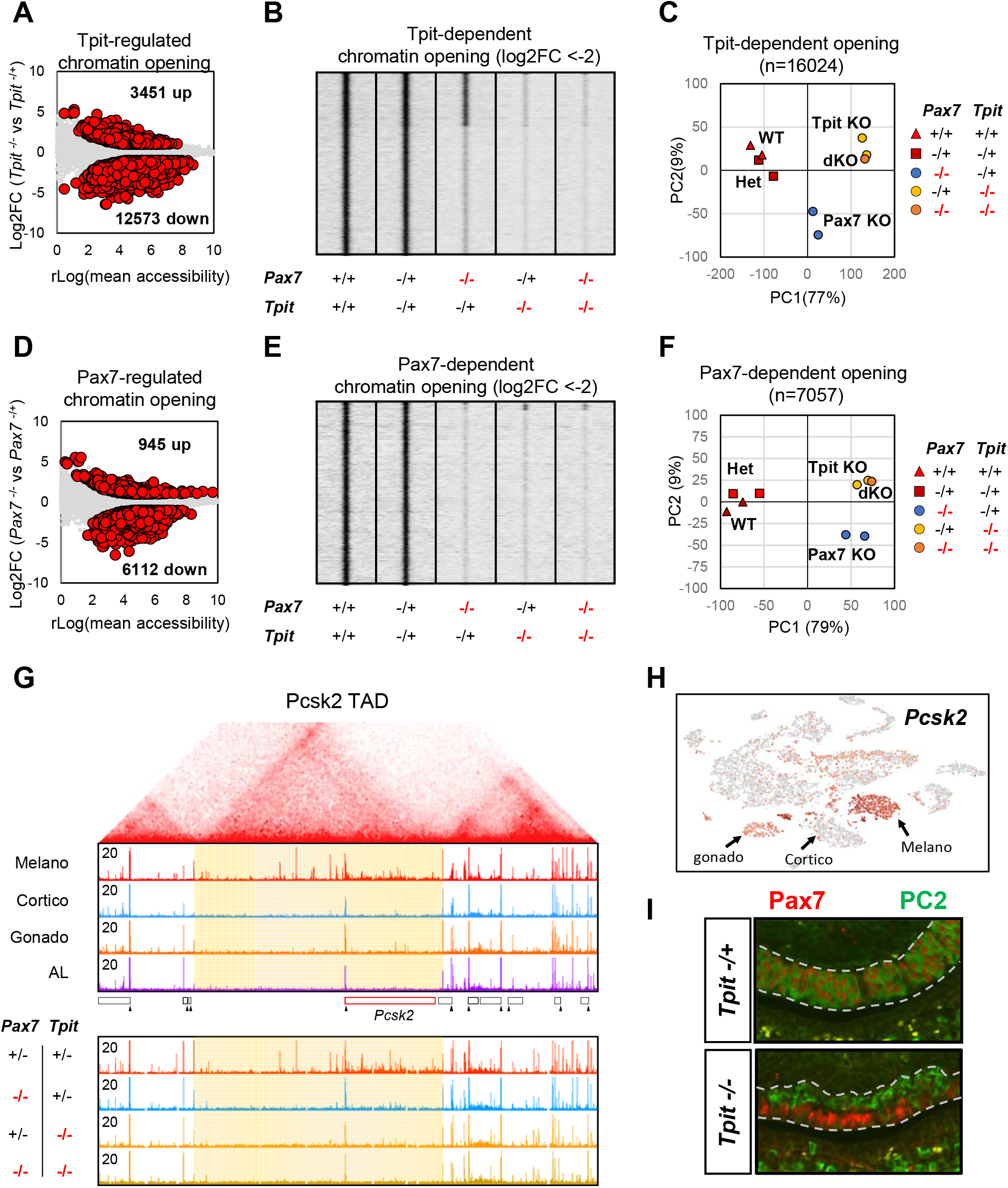
Pax7 dependent chromatin landscape requires Tpit. A. Dispersion plot showing average ATACseq rlog values (assessed by Deseq2) over the log2 fold changes of *Tpit* heterozygote versus *Tpit* knockout IL at all accessible regions. Differentially accessible regions (p-value <0.05 and log2 FC > +/− 1 as computed by Deseq2) are shown as red circles.
B. Read density heatmaps showing ATACseq signals at Tpit-dependent chromatin opening (log2 FC<−2) in the indicated mouse genotypes.
C. Principal component analysis of the ATAC signals at all 16024 Tpit regulated chromatin opening across the tested genotypes.
D. Dispersion plot showing average ATACseq rlog values (assessed by Deseq2) over the log2 fold changes of *Pax7* heterozygote versus *Pax7* knockout IL at all accessible regions. Differentially accessible regions (p value <0.05 and log2 FC > +/− 1 as computed by Deseq2) are shown as red circles.
E. Read density heatmap showing ATACseq signals at Pax7-dependent chromatin opening (log2 FC<-2) in the indicated mouse genotypes.
F. Principal component analysis of the ATAC signals at all 7058 Pax7 regulated chromatin opening across the tested genotypes.
G. Hi-C interaction map (top) from mouse ES cells^29^ around the *Pcsk2* locus showing the boundaries of the *Pcsk2* TAD. Genome browser views (bottom) of the ATACseq profiles in purified pituitary cells and ILs of the indicated genotypes at the corresponding genome location.
H. t-SNE map colored for single cell *Pcsk2* expression showing no Pcsk2 expression in corticotropes, highest expression in melanotropes and weak expression in gonadotropes.
I. Co-staining immunofluorescence for Pax7 (red) and PC2 (green) of *Tpit* heterozygote and *Tpit* knockout pituitaries from 5 days postnatal mice.

To assess the biological basis of this co-dependency, we focused on the *Pcsk2* gene, encoding the PC2 protein, a hallmark of melanotrope identity. We previously showed strong Pax7 dependency for chromatin opening of an upstream enhancer^12^ together with multiple distal elements that are ATACseq sensitive in a lineage-specific manner. Interestingly, this melanotrope-specific chromatin accessibility is confined within a topologically associated domain (TAD) and stops at the border of this TAD (Fig. 5G); similar TAD-wide opening was observed at the *Drd2* and *Grik1* loci (Supplementary Fig. 5G). In accordance with the role of Pax7 for *Pcsk2* expression in melanotropes, we found that this TAD-wide chromatin opening doesn’t occur in *Pax7−/−* animals. Consistent with our previous observation that Pax7-dependent access also depends on Tpit, chromatin opening within this TAD does not take place in *Tpit−/−* and in *Pax7−/−;Tpit−/−* animals. Thus, Tpit is required for opening of the *Pcsk2* TAD. It is noteworthy that *Pcsk2* is also expressed, albeit at low levels, in gonadotropes (Fig. 5H). This gonadotrope pattern of *Pcsk2* expression is also found in the *Tpit−/−* IL cells that have switched fate^12^; these cells occupy the dorsal side of the *Tpit−/−* mutant IL (Fig. 5I). In contrast, the ventral side of the same *Tpit−/−* IL harbors Pax7-positive cells that fail to express Pcsk2 (Fig. 5I). This confirms that Pax7 functionally requires Tpit to establish melanotrope identity.

### Productive Pax7 pioneer action in Tpit-positive cells

In order to evaluate the chromatin binding properties of Pax7 and Tpit, we compared their binding properties to those of a pituitary nonpioneer factor Pitx1 and to two pioneer factors, Neurod1 that is expressed in pituitary corticotropes^20,21^, and the reprogramming factor Sox2^6^. The heatmaps of ATACseq signals for all binding sites of these five TFs in AtT20 cells reveal striking differences between the profiles of Tpit and Pitx1, compared to those of the pioneers, Neurod1, Pax7 and Sox2 (Fig. 6A). Whereas Tpit and Pitx1 mostly (~ 95% of all sites) bind to sites that are open (ie. have a detectable ATACseq signal), a significant proportion of the pioneer binding sites have little or no ATACseq; thus for Pax7 and Neurod1, ~ 30 % of binding sites are in inaccessible chromatin whereas this is true for ~ 60% of all Sox2 binding sites. The ability to bind closed chromatin (ATAC-negative) appears to be a common property of the three pioneer factors in contrast to the non-pioneers Tpit and Pitx1.

**Fig. 6.**
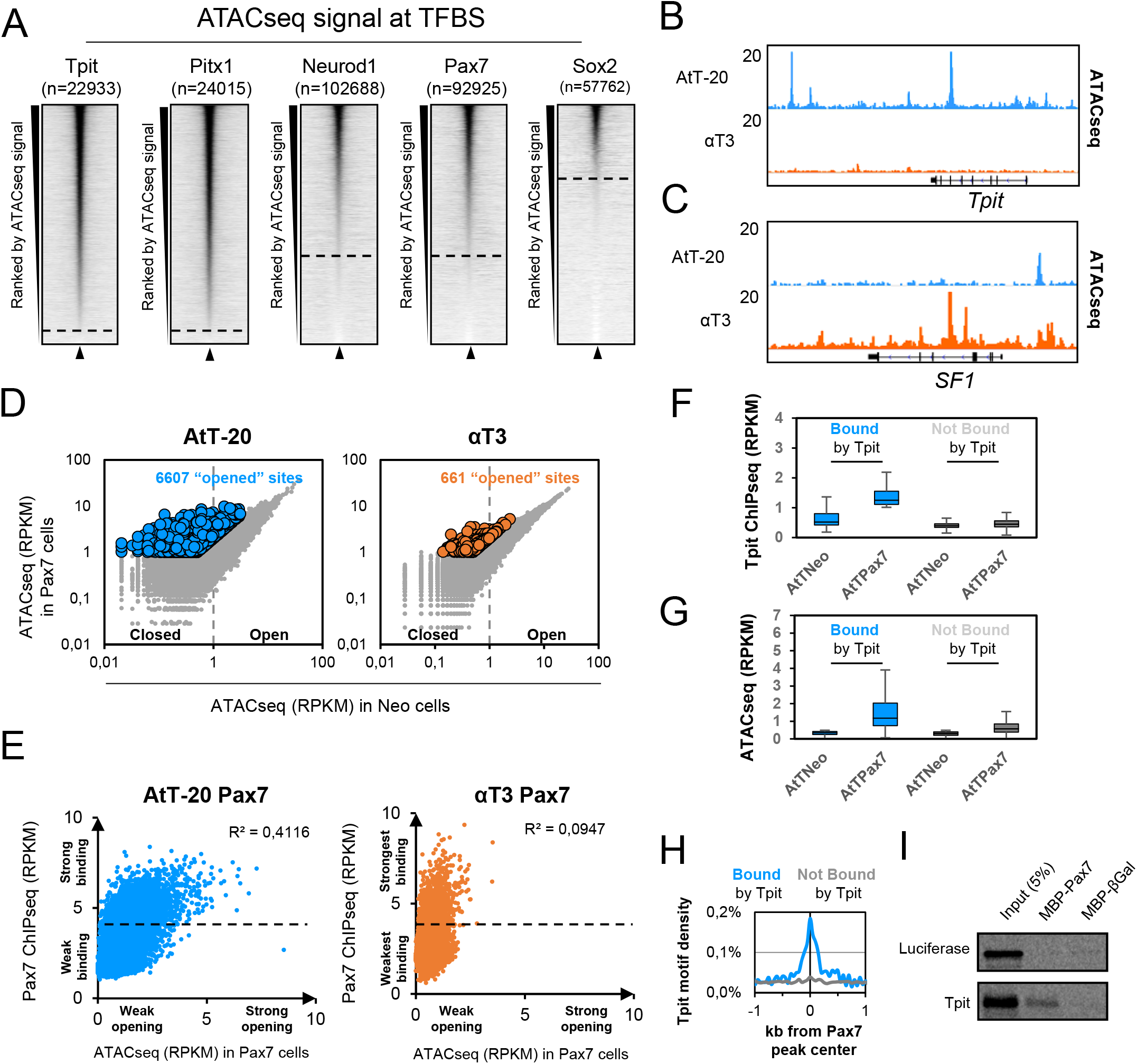
Pax7 binding on closed chromatin is only productive in Tpit-expressing cells. A. Read density heatmaps of ATACseq signal density in a 4kb window centered on binding sites for the indicated factors. The heatmaps are ranked by their decreasing ATACseq central (200bp) read densities. B,C. Genome browser views of ATACseq profiles in AtT-20 and αT3-cells at the Tpit (B) and SF1 (C) loci. D. Dispersion plots of central (200 bp) ATACseq read densities in Neo (x-axis) versus Pax7 (y-axis) expressing AtT-20 (left) and αT3 cells (right) at all Pax7 binding sites in the indicated cell lines. Colored dots represent sites with significantly stronger signals after Pax7 expression. E. Dispersion plots of ATACseq read densities in Pax7-expressing cells (x-axis) over Pax7 ChIPseq read densities (y-axis) in AtT-20 (left) and αT3 cells (right) at Pax7 binding sites with no ATACseq signal before Pax7 expression. F,G. Boxplots of Tpit ChIPseq (F) and ATACseq (G) read densities in Neo and Pax7 expressing AtT-20 cells at Pax7 sites no ATACseq signal before Pax7 expression subdivided into Tpit-bound (>1 RPKM in Pax7 expressing cells, blue) and not bound by Tpit (<1 RPKM in Pax7 expressing cells, grey). Center lines show medians; box limits indicate the twenty-fifth and seventy-fifth percentiles; whiskers extend to 1.5 times the interquartile range from the twenty-fifth to seventy-fifth percentiles. H. Tpit motif density at Pax7 sites no ATACseq signal before Pax7 expression subdivided into Tpit-bound (>1 RPKM in Pax7 expressing cells, blue) and not bound by Tpit (<1 RPKM in Pax7 expressing cells, grey). I. Pull-down assay of in vitro translated Tpit interaction with MBP-Pax7 but not with MBP-βGal.

In order to reveal the contribution of Tpit to chromatin pioneering by Pax7, we compared the impact of Pax7 expression in two different pituitary cell lines, one that expresses Tpit (AtT20) and one that does not (αT3). In agreement with expression data, the Tpit locus exhibits ATACseq signals in AtT20 cells but not in αT3 cells (Fig. 6B) whereas the locus for the gonadotrope-specific regulator SF1 presents ATACseq signals in αT3 but not in AtT20 cells (Fig. 6C).

We looked at the chromatin accessibility by ATACseq before/after Pax7 expression^12,13^ in both lineages and found that in AtT-20 cells, Pax7 binding leads to opening of 6607 regions. In contrast, only 661 sites are opened following Pax7 expression in αT3 cells and this opening is of greater magnitude in AtT-20 cells (Fig. 6D). Thus, expression of Pax7 in αT3 cells does not lead to efficient chromatin opening. We have previously identified steps of pioneering as follow: initial weak binding to closed enhancers (less than 30 minutes), stabilization of pioneer binding followed by chromatin opening^13^. Failure to perform any of these three steps should impede Pax7 pioneering. To identify which specific step limits Pax7 pioneering in αT3 cells, we compared Pax7 binding in AtT-20 and αT3 cells (Supplementary Fig. 6A). Pax7 has similar ability to bind closed chromatin in both AtT-20 and αT3 cells (Fig. 6A and Supplementary Fig. 6). We extracted all Pax7 sites bound to closed chromatin in AtT-20 and in αT3 (Fig. 6E). In both AtT-20 and αT3 cells, there are sites with strong and low binding signals. This suggest that Pax7 is able to bind strongly to closed chromatin in both cell context. However in AtT-20 cells, Pax7 binding strength correlates (r^2^=0.41) with accessibility after Pax7 expression whereas in αT3 cells, there is no correlation (r^2^=0.09) between Pax7 binding and post-Pax7 chromatin opening (Fig. 6E). We identified a subset of heterochromatin Pax7-bound sites based on their ability for Tpit binding after Pax7 expression; these sites are not bound by Tpit in absence of Pax7 (Fig. 6F) and become accessible in presence of Tpit and Pax7 (Fig. 6G). It is noteworthy that only these newly accessible sites contain the Tpit DNA binding motif (Fig. 6H). This suggests that Pax7-dependent Tpit binding also depends on the Tpit DNA motif. Notwithstanding, Pax7 and Tpit interact directly in vitro as shown in pull-down assays (Fig. 6I). The interaction between the two factors may also contribute to their cooperation. This suggests that Pax7`s ability to bind strongly to closed chromatin is not dependent on cell specific factors and that stable binding is not sufficient to drive chromatin opening. During melanotrope differentiation, this pioneer-dependent chromatin opening also requires Tpit.

## Discussion

Pioneer factors are coined as “factors that can open closed chromatin”. This label implied that pioneers were expected to directly provide this ability. The present work shows that there can be division of labor between pioneer and nonpioneer factors: in the case of Pax7 and Tpit, the former recognizes and engages pioneering sites and the latter provides the chromatin opening ability. Indeed, we show that Tpit is required for chromatin opening at sites that must first be pioneered by Pax7, and we did not find evidence for the reverse. While we envision that this is made possible by Tpit-dependent recruitment of chromatin remodeling machineries, the finding clearly circumscribes the unique features of the Pax7 pioneer function. Namely, the unique aspect of pioneer action is the ability to bind DNA sites within closed chromatin that are inaccessible to probing by techniques such as ATACseq; this appears to be a property shared with other pioneers such as Sox2, NeuroD1 (Fig. 6A) in contrast to nonpioneer factors such as Tpit or Pitx1.

For Pax7, we have shown that this initial binding to closed heterochromatin (Fig. 7A) is rapid (within 30 min, Fig. 7B) and that the first detectable change in chromatin structure at these binding sites is revealed by stabilization (within 24 h, Fig. 7C) of Pax7 binding^13^. This stabilization precedes chromatin opening (revealed by ATACseq). Here, we find that strong Pax7 binding occurs even in absence of Tpit (ie in αT3 cells, Fig. 6E): this excludes Tpit`s involvement in Pax7 stabilisation of binding. In contrast, the next step of pioneer action, namely chromatin opening, requires Tpit as it does not occur in αT3 cells. The recruitment of Tpit at a subset of Pax7 closed chromatin binding sites (Fig. 7D) that have a Tpit DNA binding site in the vicinity results in chromatin opening (Fig. 7E). It would thus be the combined interaction of Pax7 with Tpit together with the latter’s ability to bind its DNA site exposed through the initial action of Pax7 that altogether would lead to completion of pioneer action with Tpit bringing in the chromatin remodeling ability (Fig. 7E). With these interdependent functions, the cooperation between pioneer and nonpioneer factors provides robust stringency and a fail safe mechanism for triggering chromatin opening at a very specific subset of Pax7 sites.

**Fig. 7.**
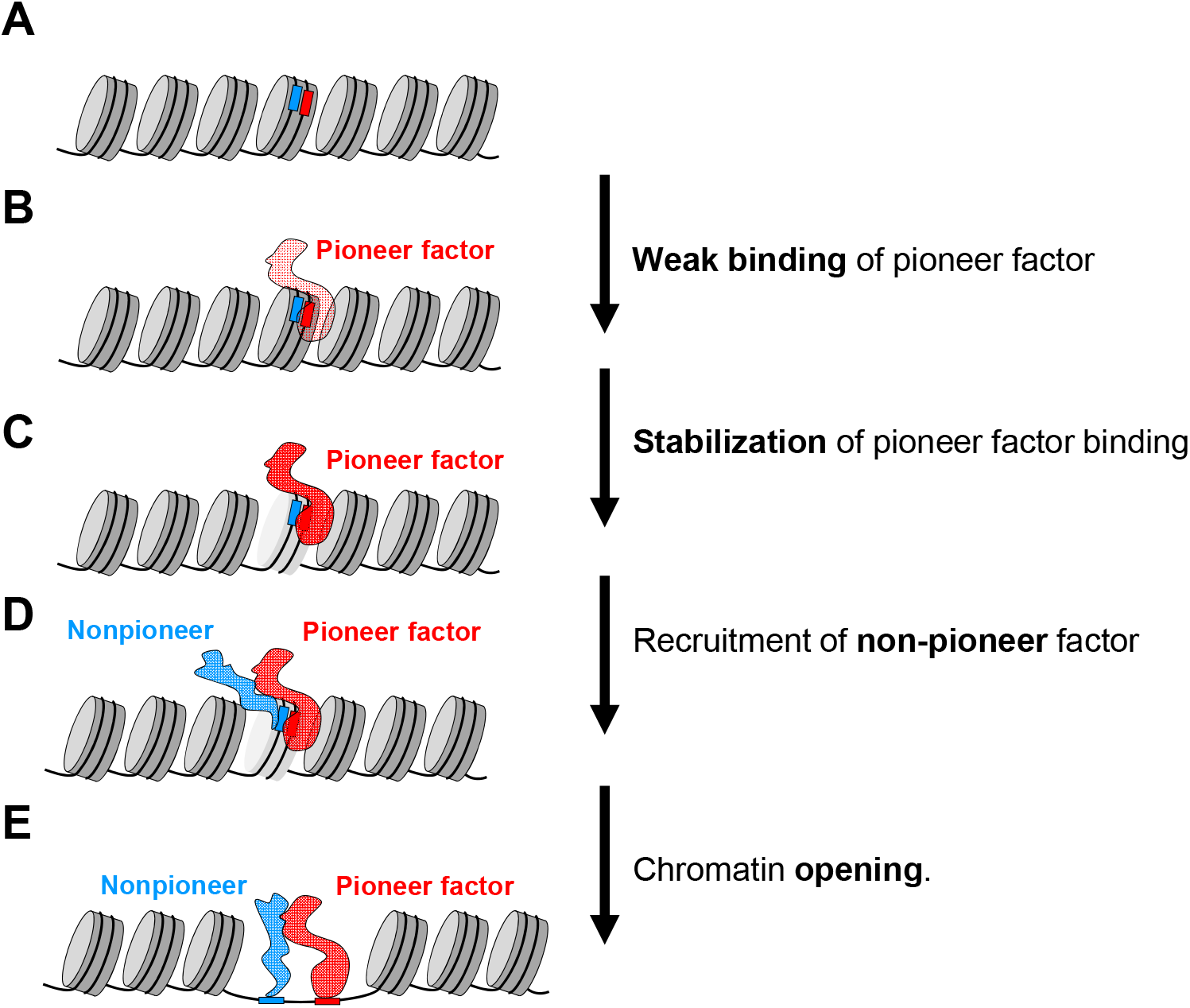
Stepwise model of pioneer and nonpioneer cooperation for chromatin opening. A. Inactive nucleosomal heterochromatin with binding sites for Pax7 (red) and Tpit (blue).
B. Weak binding of pioneer factor Pax7 to heterochromatin site.
C. Stabilized Pax7 binding with altered heterochromatin.
D. Recruitment of nonpioneer Tpit through interaction with Pax7 and its DNA binding site.
E. Accessible chromatin with both factors bound at enhancer.

The requirement for DNA binding sites for both Pax7 and Tpit at the Pax7 pioneered enhancers is reminiscent of the action of other pioneers where two different pioneers cooperate for chromatin opening. For example, the pluripotency factors cooperate with each other to facilitate the binding of other and binding of multiple pioneers is required at some sites to drive chromatin opening^7,22^. Also, there is cooperation between FoxA1 and GATA4 for chromatin binding^23^. Foxa1 also facilitates the binding of the estrogen receptor by opening chromatin^24,25^. Reciprocally, another study showed that steroid receptors can facilitate the binding of Foxa1^26^. This suggest that in the case of steroid receptors and Foxa1, both can play the role of pioneer at specific subsets of their targets. To our knowledge, the required participation of a nonpioneer for pioneer-driven chromatin opening has not been shown so far.

Prior data indicated that pioneer binding precedes gene activation, for example the binding of FoxA at liver targets precedes their activation^27^. More recently, Grainy head binding at pioneer sites was shown to determine chromatin opening (ATAC signal) but not enhancer activity^28^.

The required cooperation between pioneer and nonpioneer factors for implementation of a specific genetic program ensures specificity of action. For example, Pax7 is also involved in cell fate specification in muscle and neural tissues in addition to pituitary but its interdependent action with Tpit in the pituitary ensures that Pax7 access to pituitary-unrelated sites may not result in chromatin opening in absence of Tpit. This essential cooperation between specification and determination factors provides robustness for lineage identity by preventing mis-activation of inappropriate gene regulatory networks. This limitation to the capacity of pioneer factors would allows specific combinations of pioneers and nonpioneers to activate different regulatory networks and explain the wide variability in targets depending on the same pioneer factor in different contexts.

## ACKNOWLEDGMENTS

We are very grateful to our colleague Nicole Francis for comments on the manuscript, to Odile Neyret for NGS analyses, Eric Massicotte for FACS sorting, Dominic Fillon for microscopy, Simone Terouz for histology preparations, Dimitar Dimitrov for mouse management and to Evelyne Joyal for her expert secretarial assistance. Data analyses were possible thanks to the support of Compute Canada. This work was supported by a Foundation grant to J.D. from the Canadian Institutes of Health Research.

## AUTHOR CONTRIBUTIONS

A.M. and J.D. conceived the study, A.M., Au.B. and J.D. conceived and designed the experiments, A.M., K.S. and K.K. performed mouse tissue collection, A.M. performed ATACseq assays, A.M. and Y.G. performed ChIP experiments, K.S. performed immunofluorescence assays, J.H. did the pull-down assays, A.M. and Am.B. analyzed the single cell RNAseq, A.M. performed analysis of ATACseq and ChIPseq, A.M. and J.D. wrote the manuscript.

## Methods

### Mice, tissues, and cell culture

#### Mice genotypes and strains used in this study

Foe single cell RNAseq, a 4 month old male C57Bl/6 mouse pituitary was used; cells were dissociated as described below. For ATACseq, FACS-purified pituitary cells are isolated from 3-5 month old adult LH-cerulean^18^ or POMC EGFP^19^ C57Bl/6 transgenic pituitaries. ATACseq was performed in duplicates for each genotypes of *Pax7*^30^ and *Tpit*^9^ knockouts. Each replicate used a pool of four dissected intermediate pituitaries from 8-20 days old mice in mixed Balb/c and 129sv backgrounds. FACS analyses of intermediate and anterior pituitaries used 15-20 days-old *Pax7* knockout mice harboring the *POMC-EGFP* transgene in a mixed 129sv and Balb/c background. All animal experimentation was approved by the IRCM Animal Ethics Committee in accordance with Canadian regulations.

#### Cell lines used in this study

AtT-20 cells (obtained from the late E. Herbert in 1981 and subsequently maintained in our laboratory, with yearly negative mycoplasma tests) and αT3 cells were cultured in Dulbecco’s modified Eagle’s medium supplemented with 10% fetal bovine serum and antibiotics (penicillin/streptomycin).

#### Generation of stable Neo, Pax7 and Sox2 expressing cells

Expression vectors constructed in the pLNCX2 vector were described previously^12^. Retroviruses were packed using the EcoPack 2-293 cells (Clontech, Mountain View, CA) and infections were performed as described^12^. Selection of retrovirus-infected cell populations was achieved with 400 μg/ml Geneticin (Gibco, 11811-031). Resistant colonies were pooled to generate retrovirus-infected populations of more than 1,000 independent colonies.

### Pituitary intermediate and anterior lobe cell dissociation

After dissection, mouse pituitary intermediate or anterior lobes were dissociated as described^18^. Briefly, dissected pituitaries were separated into intermediate and anterior lobes and kept during the dissections in 300μl of dissection buffer (DMEM, 10% FBS, HEPES 10mM and DNase 100 U/ml). Anterior lobes were cut in pieces using a scalpel to facilitate tissue dissociation and digested at 37°C using 5mg/ml Trypsin for 10 minutes. We then added 2mM EDTA and incubated 5 minutes more. 10% FBS was then added to stop the dissociation and samples were centrifuged and then resuspended into 150μl of PBS 1X, 0.1% BSA, 10mM HEPES and for FACS analyses.

### 3’ end single cell RNAseq

Dissociated pituitary cells were diluted at 500 cells per μl and processed using Chromium Single Cell <u>3’</u> v2 Reagent (10x Genomics, Pleasanton, CA) following the manufacturer recommendation. Briefly, cells were passed on the channel and 9269 cells were recovered. Pituitary cells were partitioned into gel beads in emulsion for cell lysis and barcoded with oligo-dT priming and reverse transcribed. cDNA library was amplified fragmented and size selected. Samples were controlled at multiple steps during the procedure by running on BioAnalyzer. Libraries were sequenced on Hiseq 4000 with 100bp paired-end reads.

### Purification of pituitary lineages by FACS

Dissociated anterior pituitary cells from 62 LH-cerulean mice^18^ were sorted using FACSAria instrument (BD) and the gate used to define cerulean positive versus negative cells were defined by first assessing auto fluorescence of WT mice of the same strain, C57/Bl6. Cerulean positive and negative cells constituted the gonadotrope and anterior lobe (AL) samples, respectively, that were used in this study.

### FACS analyses

Dissociated anterior or intermediate pituitaries from *WT* or *Pax7-KO* mice crossed with the POMC-EGFP transgene were analysed using the FACSCalibur cell analyser (BD Bioscience). EGFP levels were quantified together with forward scatter (FSC) as an indicator of cell size and side scatter (SSC) as an indicator of granularity for both high and low EGFP-expressing cells. Experiments were repeated on 3 *Pax7* knockout and 5 wild-type litter mates in a mixed C57/Bl6 and 129sv background.

### Immunohistofluorescence

Immunohistofluorescence was performed on PFA-fixed paraffin sections as described^31^. Briefly, 5 days old *Tpit* knockout and WT pituitaries were dissected, fixed in 4% PFA, embedded in paraffin and cut into 5 μm thick sections. The following antibodies used for immunohistofluorescence: Pax7 (DSHB AB_528428), PC2 (a gift of Dr Nabil Seidah, IRCM, Montreal).

### ATACseq

All ATACseq samples were processed as previously described^13^. Briefly, 50 000 cells were washed in PBS and incubated on ice for 30 minute in a hypotonic cell lysis buffer (0.1% w/v sodium citrate tribasic dehydrate and 0.1% v/v Triton X100) and centrifuged (5 minutes at 2000g at 4°C). Cells were then incubated 30 minutes on ice in cell lysis buffer (10mM Tris-HCl, pH7.4, 10mM NaCl, 3mM MgCl2, 0.1% v/v IGEPAL CA-630. After centrifugation (5 minutes at 2000g at 4°C), the nuclei pellets were resuspended in transposase Master Mix (1.25 μl 10x TD buffer, 5 μl H2_O_ and 6.5 μl of Tn5: Illumina Nextera Kit; FC-121-1031) and incubated for 30 minutes at 37°C. Samples were purified using the DCC purification columns (Zymo). The eluted DNA was barcoded for multiplexing of samples using Nextera barcodes and PCR-enriched using the Phusion kit. Libraries were recovered with GeneRead Purification columns. Samples were then evaluated by qPCR to test enrichments and sequenced on Illumina Hiseq 2500 with 50bp or 125bp paired-end reads according to Illumina’s recommendation.

### ChIPseq

ChIPseq were performed as previously described^32^. At least 3 immuno-precipitations were pooled per ChIP experiments. Library and flow cells were prepared by the IRCM Molecular Biology Core Facility according to Illumina’s recommendations and sequenced on Illumina Hiseq 2500. The following antibodies were used for ChIPseq: FlagM2 (Sigma F3165), Neurod1^33^, Sox2 (Ab59776, Abcam).

### Pull-down assay

MBP fusion proteins coupled to maltose amylose beads were produced as described^34^. ^35^S labeled proteins were synthesized *in vitro* using the TNT T7 Quick for PCR DNA kit (Promega, L5540). Labeled proteins were incubated with MBP-tagged proteins in TNEN50 (50mM Tris pH7.5, 5mM EDTA, 50mM Nacl, 0.1% NP-40) with 1mM PMSF and 2% BSA for 4 hours at 4°C. Beads were washed three times with 1ml TNEN_125_. Bound proteins were resolved by SDS-Page and visualized by autoradiography.

### Data Analyses

#### Single Cell RNAseq analyses

Using Cell Ranger v2.1.1 (10X Genomics), reads were aligned on the mm10 mouse reference genome using the default parameters of Cell Ranger to generate unique molecular identifier counts for each genes across the 9269 cells that were profiled. We obtained an average of 25220 reads per cell and we detected 1807 median genes per cell. Using the Cell Ranger pipeline with default parameters, we generated a gene-barcode matrix, principal component analysis and dimensionality reduction using the t-SNE algorithm. Unbiased clustering of single cells was performed using Cell Ranger which combines K-means clustering and graph-based clustering uncovering 12 clusters. We used Loupe Cell Browser (10X Genomics) to visualize the t-SNE plot with colored cell according to their assigned cluster or colored by gene UMI. Loupe Cell Browser was also used to perform local differential expression analyses of clusters 1, 2, 4, 5, 7, 8. Cluster corresponds to 4 islands of cells that each express a different hormone gene, Gh, Prolactin, Lh and POMC. Differential analysis of the four sub-clusters against their matching hormone-expressing cell cluster showed that mitochondrial RNAs are down-regulated in each case and this is an indicator of low quality cells^16^. Thus to avoid confounding effects, cluster 3 was not included in all further analyses.

#### ATACseq

Reads were trimmed, if required, to obtain a read length of 50bp and aligned to the mm10 mouse reference genome using Bowtie v2.3.1^35^ with the following parameters: --fr --no-mixed --no-unal. Sam files were converted into tag directories using HOMER v4.9.1^36^ and into bam files using Samtools v1.4.1^37^ view function. Tag directories were used to generate the normalized BigWig files with Homer using the command makeUCSCfile with the parameters: -fsize 1e20 -res 5 -fragLength 100. Peaks were identified by comparing each sample replicate to sequenced input DNA from pituitary using MACS v2.1.1.20160309^38^ callpeak function using the parameters: -f BAMPE --bw 250 -g mm --mfold 10 30 -p 1e-5. Peaks with an associated pvalue less than 10^−5^ were kept. First we compared ATACseq profiles of purified pituitary cells: melanotrope (2 replicates), corticotropes (2 replicates) gonadotropes (1 replicate) and whole AL (1 replicate). Peaks from all datasets from purified pituitary cells were merge using HOMER v4.9.1 mergePeaks tool to obtain a file with all unique positions from the ATACseq datasets. This list was clustered by k-means in 2 clusters for each samples giving the 16 combinations of ATAC clustering as represented in a heatmap in Fig. 2B. Peaks from all datasets from the various genotypes of *Pax7* and *Tpit* knockout ILs were merged together using HOMER v4.9.1 mergePeaks tool to obtain a file with all unique IL positions from all ATACseq datasets. ATACseq signals were quantified in these different datasets using the analyzeRepeats.pl HOMER command and differential accessibility analyses was performed using getDiffExpression.pl with default parameters which uses Deseq2. Peaks showing a differential p-value less than 0.05 and a fold change of 2 fold or more were considered differentially accessible.

#### ChIPseq

We mapped ChIPseq reads on the mouse genome assembly mm10 by using Bowtie v1.1.2 with the following settings: bowtie -t -p 4 --trim5 1 --best mm10 –S. Sam files were converted into tag directories using HOMER v4.9.1 and into bam files using Samtools v1.4.1 view function. Peaks were identified by comparing each sample to its control (IP Flag for Pax7, IP IgG for others) using MACS v2.1.1.20160309 callpeak function using the parameters: --bw 250 -g mm --mfold 10 30 -p 1e-5. Peaks with an associated p value less than 10^−5^ were kept.

#### Data presentation

Heatmaps and average profiles were generated using Easeq^39^, We used IGV^40^ to visualize the BigWig files on the genome. Principal component analysis, clustered Heatmap associated with dendrograms from Figures 1D,5C,F were generated using ClustVis^41^.

## Legends to Supplementary Figures

**Supplementary Fig. 1.**
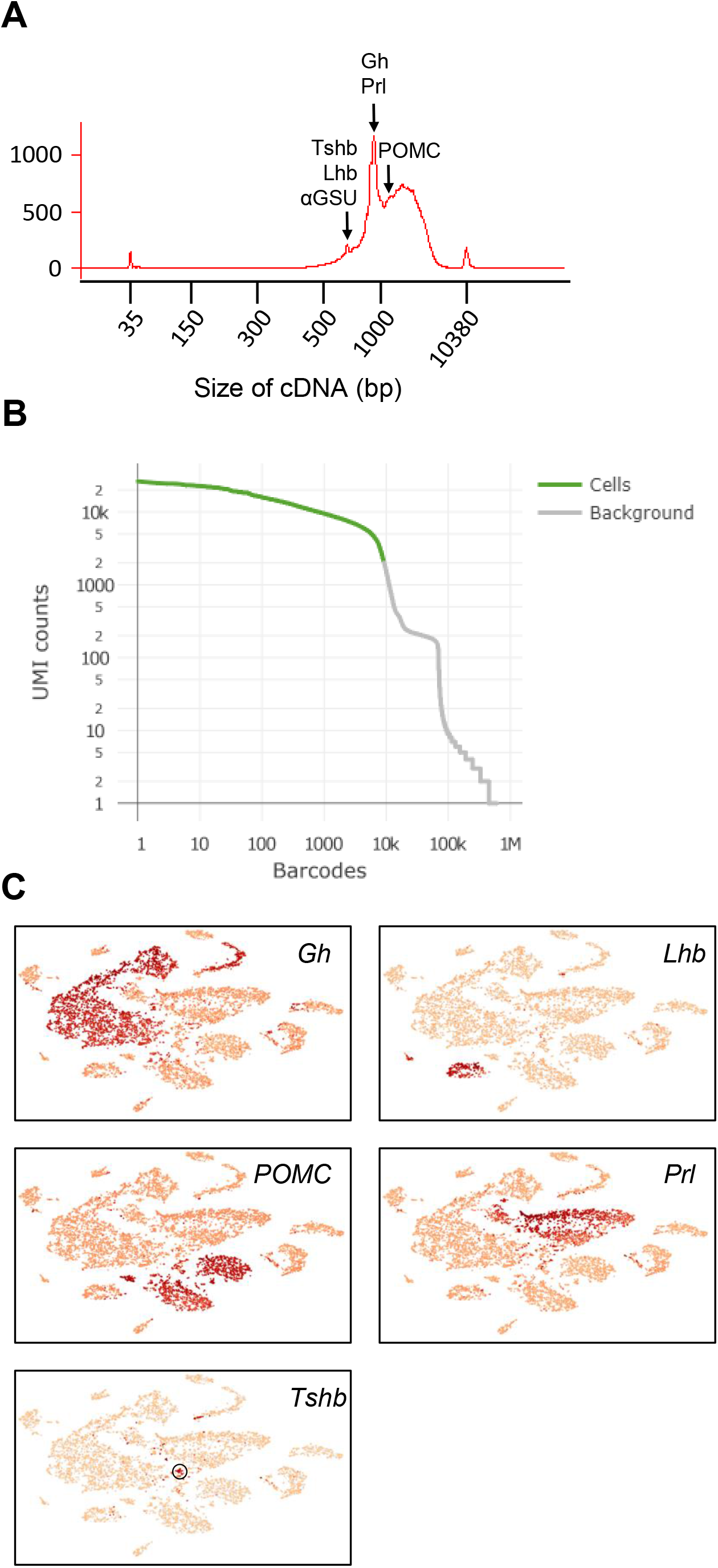
Validation of single cell RNAseq data. (A) Bioanalyzer profile showing size distribution of cDNAs from dissected pituitary cells. (B) Single cell RNAseq plot showing barcode numbers (x-axis) over the number of UMI per barcode (y-axis). The threshold (colored) used for selecting the 9269 cells analysed. (C) t-SNE map showing color-coded expression of hormone genes Gh, Lhb, POMC, Prl and Tshb.

**Supplementary Fig. 2.**
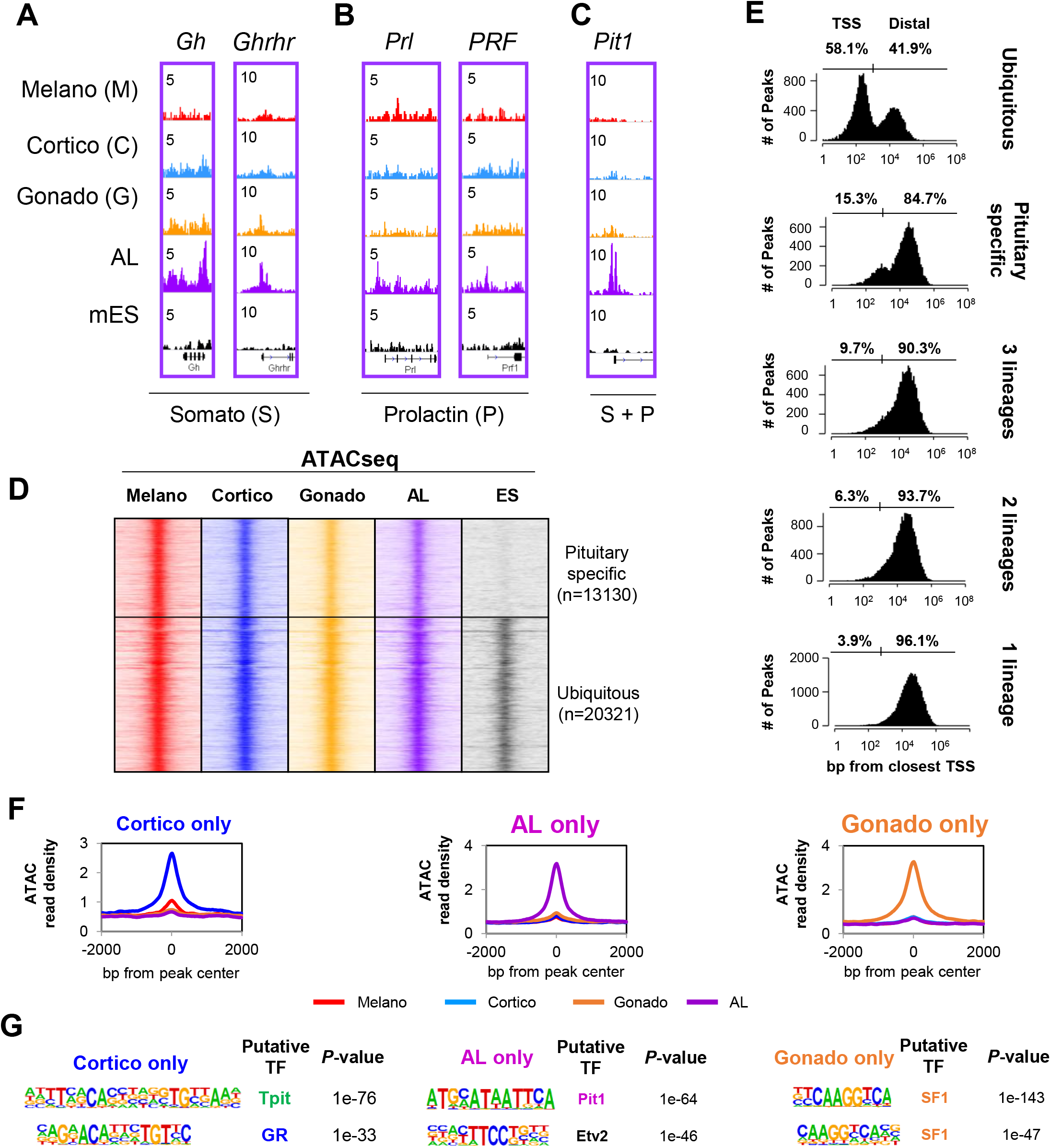
Chromatin landscapes in other pituitary lineages. (A-C) Genome browser views (IGV) of ATACseq profiles from the indicated pituitary lineages at genes marking the identity of somatotropes (A), lactotropes (B) and both (C). (D) Read density heatmaps showing ATACseq signals at sites of accessibility common to all four pituitary lineages clustered by ATACseq signals in mouse embryonic stem cells (GSE64058). (E) Genomic distribution of the distances between the indicated category of ATACseq peaks and the closest TSS. (F) Average profiles of ATACseq signals for the four indicated lineages at cortico-(blue), anterior lobe (purple) and gonado-(orange) specific ATACseq sites. (G) Motif enrichments (assessed by HOMER) under the indicated category of ATACseq peaks.

**Supplementary Fig. 3.**
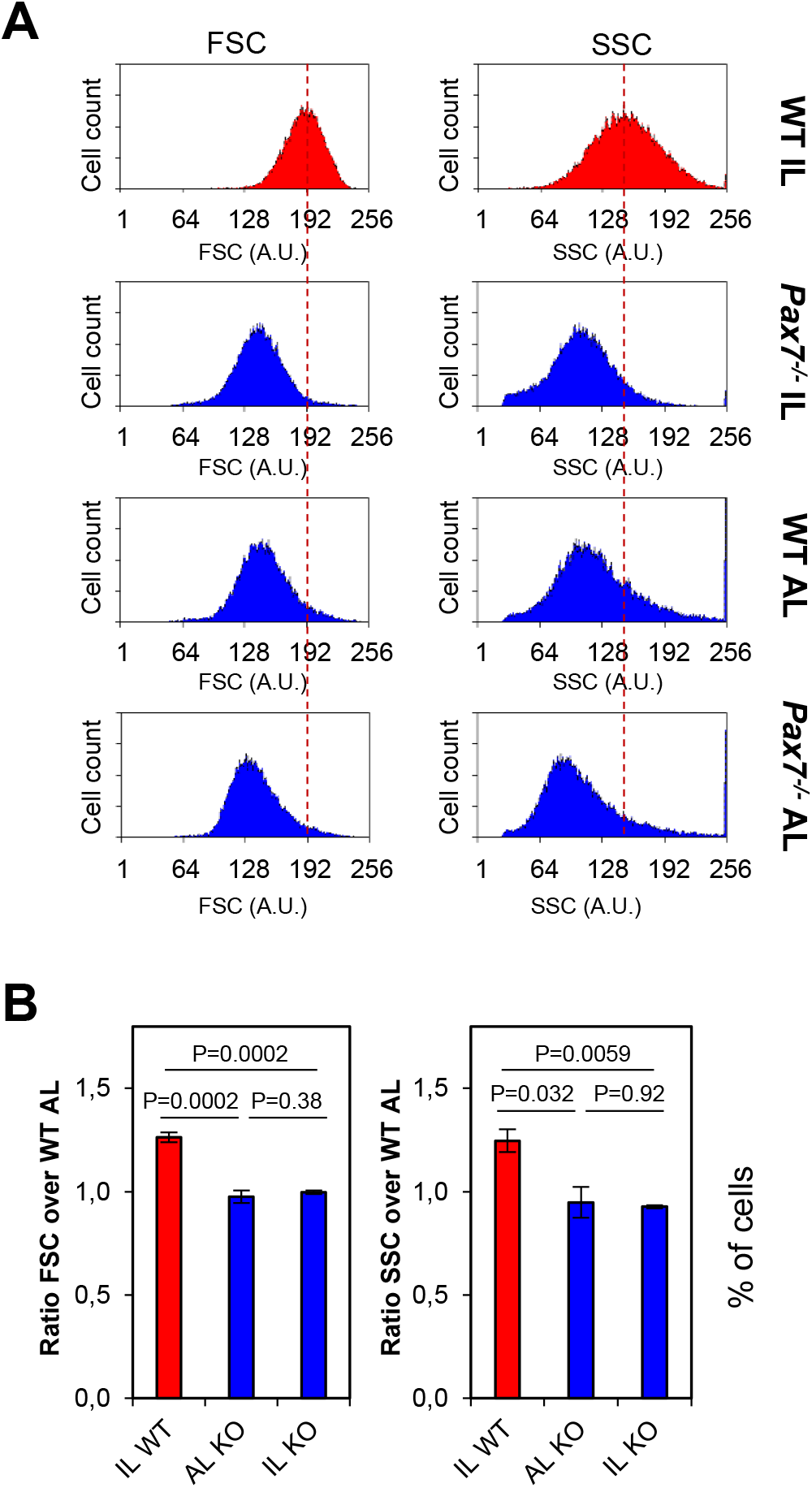
FACS analyses of wild-type (WT) and *Pax7−/− POMC-EGFP* pituitary cells. (A) Representative FACS profiles showing forward (left) and side (right) scatter for cells from wild type and *Pax7^−/−^* IL and AL. (B) Bar graphs showing the ratios of forward (right) and side (right) scatter for EGFP positive cells from wild type (n=5) and *Pax7^−/−^* (n=3) IL and AL.

**Supplementary Fig. 4.**
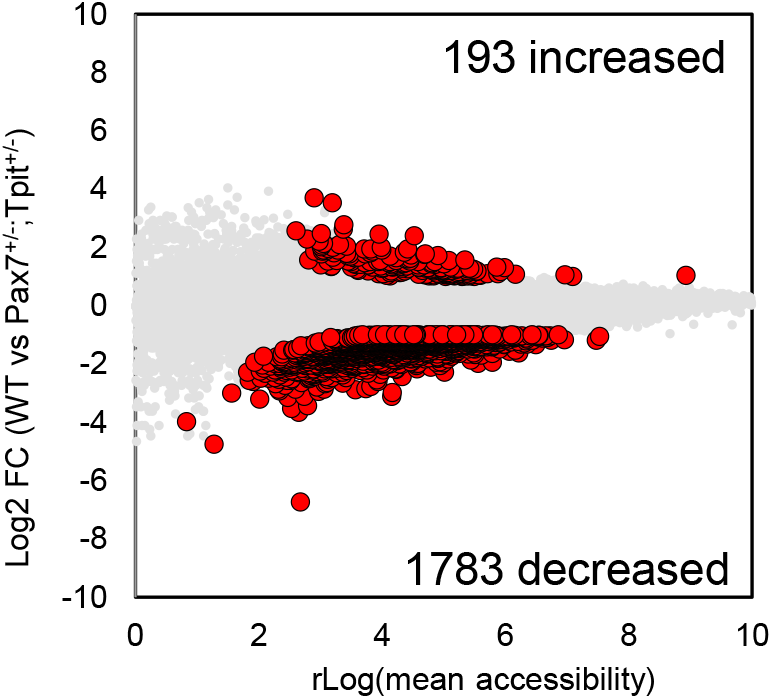
Dispersion plot comparing ATACseq signals in WT and Pax7^+/−^;Tpit^+/−^ IL cells. Dispersion plot showing rLog values of accessibility (ATACseq, x-axis) in wild type and *Pax7*^+/−^; *Tpit*^+/−^ IL over log2 fold changes in wild type versus *Pax7*^+/−^; *Tpit*^+/−^ IL. Red circles identify the differentially accessible regions (p value<0.05, Log2 FC >+/−1).

**Supplementary Fig. 5.**
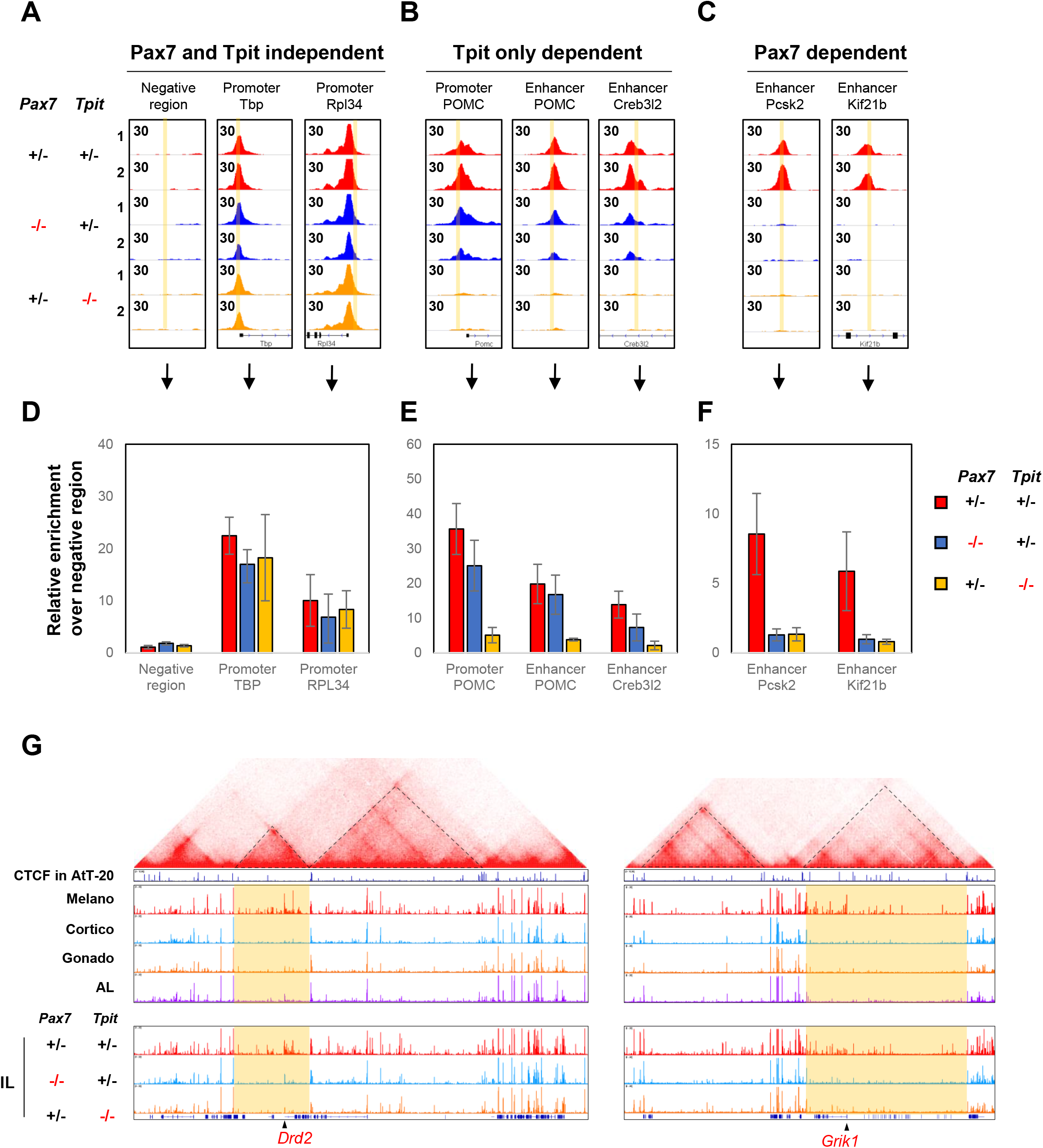
Q-PCR validation of Pax7 and/or Tpit-dependent accessibility (ATACseq). (A-C) Genome browser views (IGV) of ATACseq profiles from the indicated genotypes at unaffected (A), Tpit-only dependent (B) and Pax7 dependent (C) sites. Regions amplified in the qPCR measurements of Figure S5D-F are highlighted in yellow. (D-F) Relative enrichments over a negative region, measured by qPCR of ATACseq libraries at unaffected (D), Tpit-only dependent (E) and Pax7 dependent (F) sites. (G) Hi-C interaction map (top) from mouse ES cells (Bonev et al., 2017) around the *Drd2 (left)* and *Grik1 (right)* loci showing the boundaries of their respective TADs. Genome browser views (bottom) of the ATACseq profiles in purified pituitary cells and ILs of the indicated genotypes at the corresponding genome location.

**Supplementary Fig. 6.**
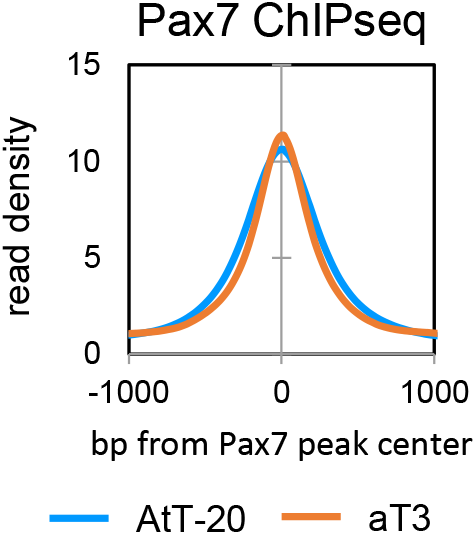
Comparison of Pax7 ChIPseq data in AtT-20 and αT3 cells. Average profiles of Pax7 ChIPseq in AtT-20 (blue) and αT3 (orange) cells at the best 2000 Pax7 peaks from each lineage showing similar enrichments in both ChIPseqs.

