## Supplemental Table 1 for "Pioneer and nonpioneer factor cooperation drives lineage specific chromatin opening"

| | Forward | reverse |
|---|---|---|
| Negative region | GTAGGAGGTACCAATAACTGTG | TAGGACCAAGAAGATGACTCT |
| Negative region 2 | CCAGAGCATTGGCTATGTAA | TGCCCTAAGATGCAGTAAAG |
| Promoter Tbp | GTAGAACGCTTGCTTGGGCTTGAT | AGCTGTGAGACACTGGGAAGGAAA |
| Promoter RPL34 | AAGGGCCACGATGCCTTTAT | AAGTGTGTGCTCTGGCTGAA |
| Promoter POMC | TGGTTTCACAAGATATCACACTTTCCC | TCGGAGTGGAATTACCTATGTGCG |
| Enhancer POMC | ATCTTCATGTGTCCGGTGTTGGGT | ACCAGGGTGTGAGTACGCTACAAA |
| Enhancer Creb3l2 | ACCAGAGTTCTGCCCTTTGA | TGTCACCAAGTGCTACTCCA |
| Enhancer Pcsk2 | GAATTCTATGCCATCAGTAGC | ACACATCCCTCTCAATAAGT |
| Enhancer Kif21b | ACTGGTAGCACCTGGTCTGAAT | TGCTACTCACTCCAGCCAACTT |
